# Diversity and relative abundance of Collembola in a wheat (*Triticum aestivum*) field at Aligarh

**DOI:** 10.1101/580811

**Authors:** Mohammad Jalaluddin Abbas, Hina Parwez

## Abstract

Collembolans are novel indicators of soil quality as they are enormously diversified in agricultural soils. However, their abundance is ever dynamic due to the dynamic climatic conditions. In order to ascertain the diversity and relative abundance of Collembola associated with wheat field, soil samples were taken at weekly intervals from selected field of Quarsi village located at Aligarh district of Uttar Pradesh (India). Total 20 samples were taken for the site study during the investigation period and soil microarthropods extracted by using modified Tullgren funnel apparatus. The results of our study showed that, the species diversity of Collembolans mainly consists of individuals belonging to family Entomobryoidae, Isotomidae, Hypogastruridae and Sminthuridae. Among these, Hypogastrurides were dominant (56.84 %) in entire community of Collembola. Soil temperature was negatively correlated (r = −0.932, P<0.05) with reference to Collembolans population, whereas soil moisture (r = 0.502, P>0.05) as well as available nitrogen (r = 0.656, P>0.05) both were positively correlated. The highest population of Collembolans was recorded at neutral pH level. In terms of numbers of soil microarthropods, Collembolans apparently constituted a better population than the other diverse group of soil microarthropods such as Acari(mites). The present study has shown profound diversity of Collembolans and highlights the significance of the variety of chemical and edaphic factors which regulate the fluctuation and diversity of microarthropods in a varied manner.

## Introduction

In an agro based country like India, the importance of soil ecosystem in terms of crop productivity cannot be over emphasized (Parwez and Sharma-2004). The functions for crop productivity are directly or indirectly linked with various functional attributes (biotic & abiotic) in an ecosystem. Thus, soil ecosystem is a medium by which both below ground as well as above ground living beings are benefited. Still, soil ecosystem is an important yet complex medium that play a potential role in maintaining the biological diversity in below ground and crop productivity for above ground. The productivity of soil directly or indirectly depends on the activities and interactions of soil micro fauna and micro flora. The presence and absence of certain micro flora had an influence over the population of very diverse soil microorganisms such as microarthropods. The grazing activity of soil microorganisms is a key function by which they facilitate the soil environment qualitative. Among the soil microarthropods, the interaction of Collembolans population with that of habitats has its profound affect on the nature and fertility of soil. Thus, an eco-friendly functional approach of Collembola has generalized and their diversity plays a very crucial role in an agro-ecosystem specifically in wheat field.

Many research workers have concluded that, Collembolans are good bio-indicators of arable soil quality (Rusek-1998), are very sensitive to variations in soil environment (Parisi-2001) and conditioning to detritus for microbial breakdown as well as farming soil to form soil micro-structure (Chahartaghi-2005). Thus, we focused only on Collembolans in this study. The reason for this selective field study is much important that, wheat is an important crop and used more than 70% human beings as for food supplement in India. Still, wheat is a first line crop in Uttar Pradesh; however, its medium production recorded in this region from last two decades.

There is a paucity of information on soil microarthropods and their population dynamics in agricultural soils of this region in relation to the diversity of Collembolans or other soil micro arthropods. However, some other comparative studies reported from West Bengal (Chaudhury and Roy-1967, 1971, 1972, Hazra and Chaudhury-1990, Chaudhury et al. 1978, Hazra-1978a and 1978b and Mitra et al. 1977, 1981, 1983). In the present study, we investigate the diversity and relative abundance of Collembolans from wheat field at Aligarh. The parameters considered in our study are density, relative-abundance, fractional population, absolute frequency and diversity along with concomitant changes with a variety of physical and chemical factors.

## Materials and Methods

### Area of study

The area selected for study is situated at Aligarh. It is a flat topographical area, located in western part of UP at latitude 27-54’N, longitude 78-05E’ and altitude 187.45 meter above sea level. It is a subtropical zone with fluctuating climatic conditions consisting of four different seasons characterized by extreme winter and summer followed by medium to heavy rainfall during monsoon months and a post monsoon sweet spring. In hot dry summer, the temperature rises up to 48 ºC, while in winter-cold, the temperature down up to 2 ºC. Relative humidity also fluctuates with the sudden change of environmental temperature and with rainfall patterns. Such widely varying climatic conditions provide a variety of ecological niche to soil dwelling organisms and interesting for soil ecological studies in this region.

### Study site

A site was selected at Quarsi village that is situated at the outskirts of Aligarh city. The crop that we consider was winter wheat (*Triticum aestivum*) that was grown in the very beginning of November. The site was well managed by its farmer in terms of ploughing, irrigation, and fertilizers used time to time in the field. The organic fertilizer used by its farmer in the field ones before 15 days of growing time of wheat. The chemical fertilizer used in the field was die ammonium phosphate (DAP), ones after the growing period at about one month. The field was irrigated ones before the growing of wheat at about 20 days and four times after the growing with every one month interval. The soil of site was alluvial type, a mixture of sand, silt and clay. The approximate area of the sampling site was two acre. Samples were taken regularly at the rate of 4 samples per month and from random points inside the field.

### Sampling, extraction and Identification of soil microarthropods

As mentioned earlier, samples were taken every week regularly and the points selected with in the site were distributed randomly. Total 20 samples were taken for the site study during five month of wheat crop. Modified Tullgren funnel apparatus was used for the extraction of soil microarthropods. The power of bulbs used was 60 watts. The size of corer varies with the amount of soil examined. All microarthropods were collected in collecting vials which contained 70 % alcohol with few drops of glycerol and they separated and mounted with DPX. A stereo-zoom-microscope was used to identify the soil microarthropods up to the level of their order or, family using a range of taxonomic keys (O’ Connell and Bolger 1997).

### Statistical Analysis

To study the population dynamics of Collembola, the parameters considered are density, abundance, fractional population, absolute frequency and relative abundance etc. One way analysis of variance (ANOVA) was calculated for the significance (P> 0.05) of population of collembolans with edaphic factors (Prasad 2007). Relative abundance (%) calculated by following formula-

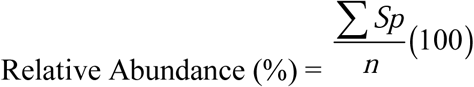

Where, *S*_*p*_ = population of particular group/species within time selected, and

*n* = total number of individuals present

## Results

An unexpected abundance and diversity of Collembola found during the investigation period. The species richness of soil Collembolans was low while the individual population was high in most of sampling cases. The diversity indices such as density and abundance of Collembola population were higher (51.50) in January simultaneously than compare to other months (table-2). Both, Poduromorpha (56.84 %) and Entomobryomorpha (36.91 %) constituted a better understanding than Symphypleona (6.24 %), (table-1). Poduromorpha always constituted a better population (density as well as abundance) than compare to others. March was the least diversified month that may be due to the increased environmental temperature and decreased soil moisture content in soil (see table-2 and figure-1). Collembola Absolutely frequent up to 88.35 % with an average of 72.62 % during the investigation period (table-2). The average density and abundance of Collembola found up to 28.25 and 33.20 during investigation period (table-2) and these values found more than expected for the site study.

**Figure: (1).**
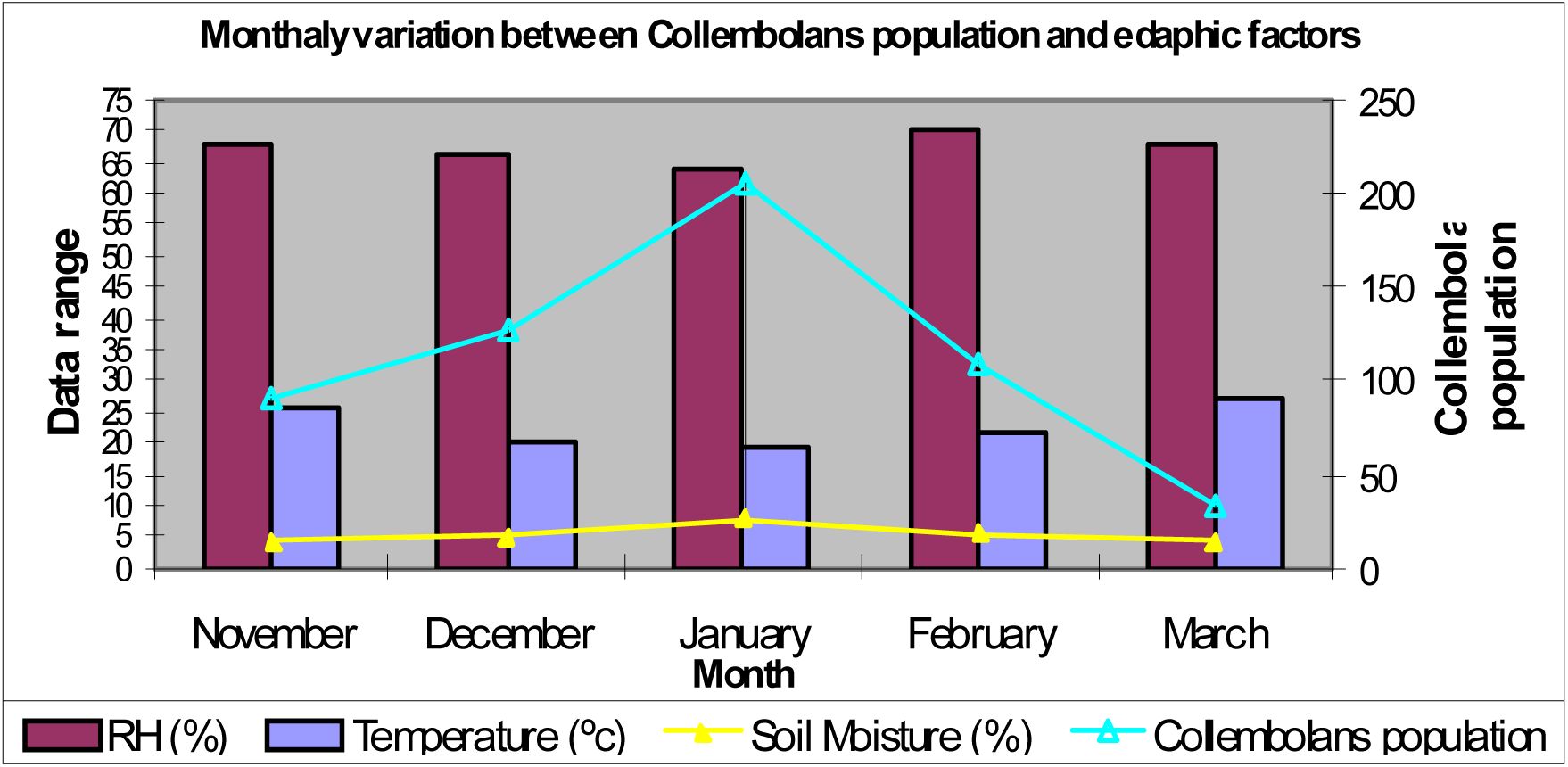
Monthly variation between Collembola population and edaphic factors.

**Table: (1).** Relative abundance (%) of Collembola communities in a wheat field at Aligarh.

|  | **Poduromorpha** | **Entomobryomorpha** |  | **Symphyleona** |
| --- | --- | --- | --- | --- |
|  | (Hypogastruridae) | Entomobryoidae | Isotomidae | (Sminthuridae) |
| November | 56.04 | 18.68 | 19.78 | 5.49 |
| December | 50.79 | 26.98 | 16.67 | 5.56 |
| January | 61.65 | 18.93 | 14.08 | 5.34 |
| February | 56.36 | 23.64 | 14.55 | 5.45 |
| March | 59.37 | 15.63 | 15.63 | 9.37 |
| *Average* | *56.84* | *20.77* | *16.14* | *6.24* |

**Table: (2).**
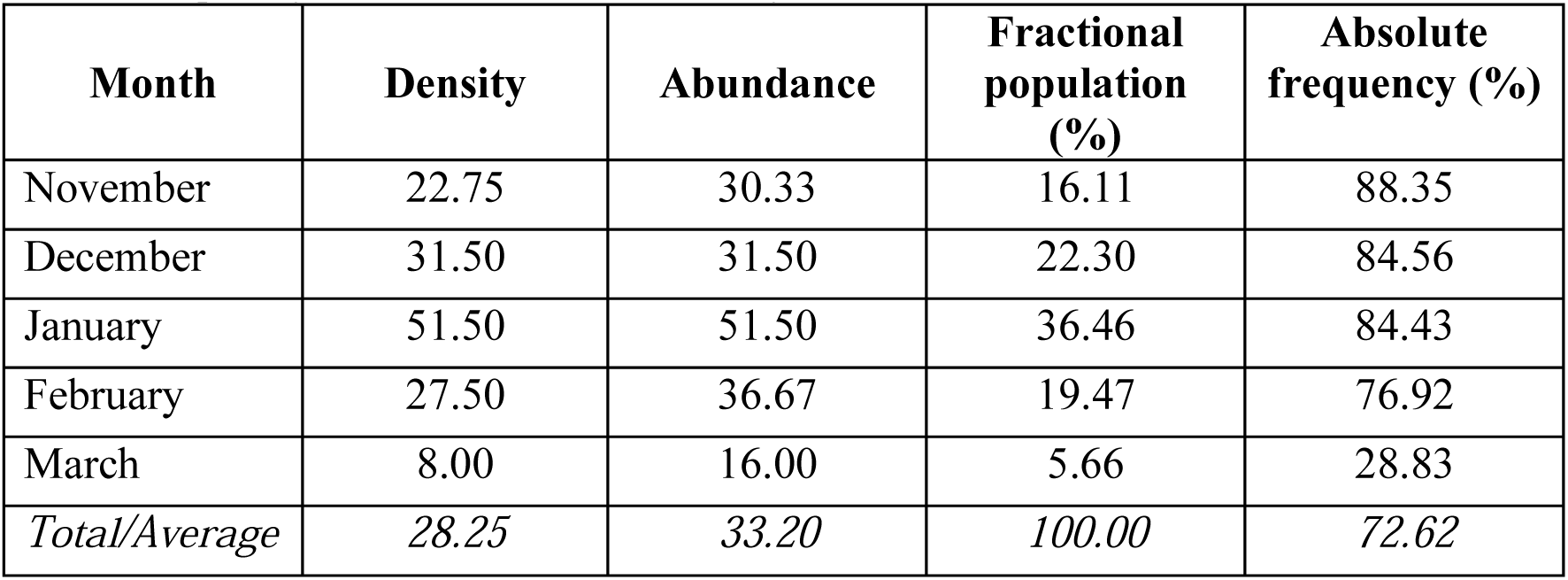
Monthly variation between density, abundance, fractional population and absolute frequency of Collembola community.

| **Month** | **Density** | **Abundance** | **Fractional population (%)** | **Absolute frequency (%)** |
| --- | --- | --- | --- | --- |
| November | 22.75 | 30.33 | 16.11 | 88.35 |
| December | 31.50 | 31.50 | 22.30 | 84.56 |
| January | 51.50 | 51.50 | 36.46 | 84.43 |
| February | 27.50 | 36.67 | 19.47 | 76.92 |
| March | 8.00 | 16.00 | 5.66 | 28.83 |
| *Total/Average* | *28.25* | *33.20* | *100.00* | *72.62* |

Soil carbon content and available nitrogen ranges between 0.21-0.25 % and 253-281ppm (figure-2). Soil temperature and relative humidity both were ranged between 18.3 to 27.4ºC and 63.5 to 70.0 % during investigation period (figure-1). It was reported that the increased population of Collembolans may be due to increased carbon content in the soil (figure-2). This study revealed that the species diversity of Collembola correspond to Entomobryoidae, Isotomidae, Hypogastruridae and Sminthuridae. Among these, Hypogastrurides were dominant (56.84 %) in entire community of Collembola (table-1). Temperature was negatively correlated with reference to Collembolans population (r = −0.932, P<0.05), whereas soil moisture (r = 0.502, P>0.05) as well as available soil nitrogen (r = 0.656, P>0.05) both were positively correlated. Soil organic carbon contents vary significantly with the population fluctuation of collembolans (figure-2).

In the present study, the density of Collembolans found between 1019-6560/m² in crop selected (*Triticum aestivum*) that was low to medium in this region of agricultural soil. Both, acidic and basic natures of soil were recorded as the effective ranges for the diversity of soil microarthropods specifically for collembolans population because, the higher population of Collembolans recorded at neutral pH level. In terms of numbers of soil microarthropods, Collembola apparently constituted a better population than the other diverse group of soil microarthropods such as Acarina (mites); however they are not considered in this study.

## Discussion

It is well known that Collembolans are the important functional agents of soil ecosystem specifically in agricultural lands because they provide an eco-friendly functional approach directly or indirectly for plant growth by help in recycling of nutrients, decomposition of organic matter in soil, to support in formation of soil micro-structure and host for many pathogens such as fungi, bacteria, nematodes and protozoans etc. Thus, rich diversity of Collembola can be the indicator of soil health status because of their very specific habitats and contribution to ecological functional medium in soil ecosystem. Some of them are excellent indicators of biotic and abiotic disturbances in agricultural ecosystem. Bengtsson (1998) also proposed that, studies should be focused on key tone species and functional groups rather than of species diversity. Thus, we focused only on Collembolans in our study that is a very specific functional group including their whole diversity. It may offer the interest of such natural resource conservation as well as long term functioning of native agricultural soils.

As shown in the figure 2, the soil organic carbon content was positively correlated with reference to collembolans population. This may be due to the proper use of organic fertilizer by the farmer in the field. Thus, it can be stated that soil organic carbon contents vary significantly with the population fluctuation of collembolans. The level of the available nitrogen increased in the soil continuously throughout the investigation period (figure-2) that was due to the use of DAP fertilizer by farmer, after that the nitrogen released through the leaching of ammonium into the soil. The increased level of available nitrogen in soil does not affect the collembolans population. However, the medium concentration of available nitrogen in soil was favorable for higher diversity of collembolans (figure-2). The seasonal fluctuation in soil temperature may be another reason for the population dynamics of collembolans. It was observed that the cool temperature was favorable for high diversity of collembolans because from the study observations, we found that the rising temperature negatively correlated to the collembolans population (figure-1). The fluctuations in the soil moisture regimes can be the possible factor for the population dynamics of collembolans during study time that was positively correlated to the population of collembolans (figure-1).

**Figure: (2).**
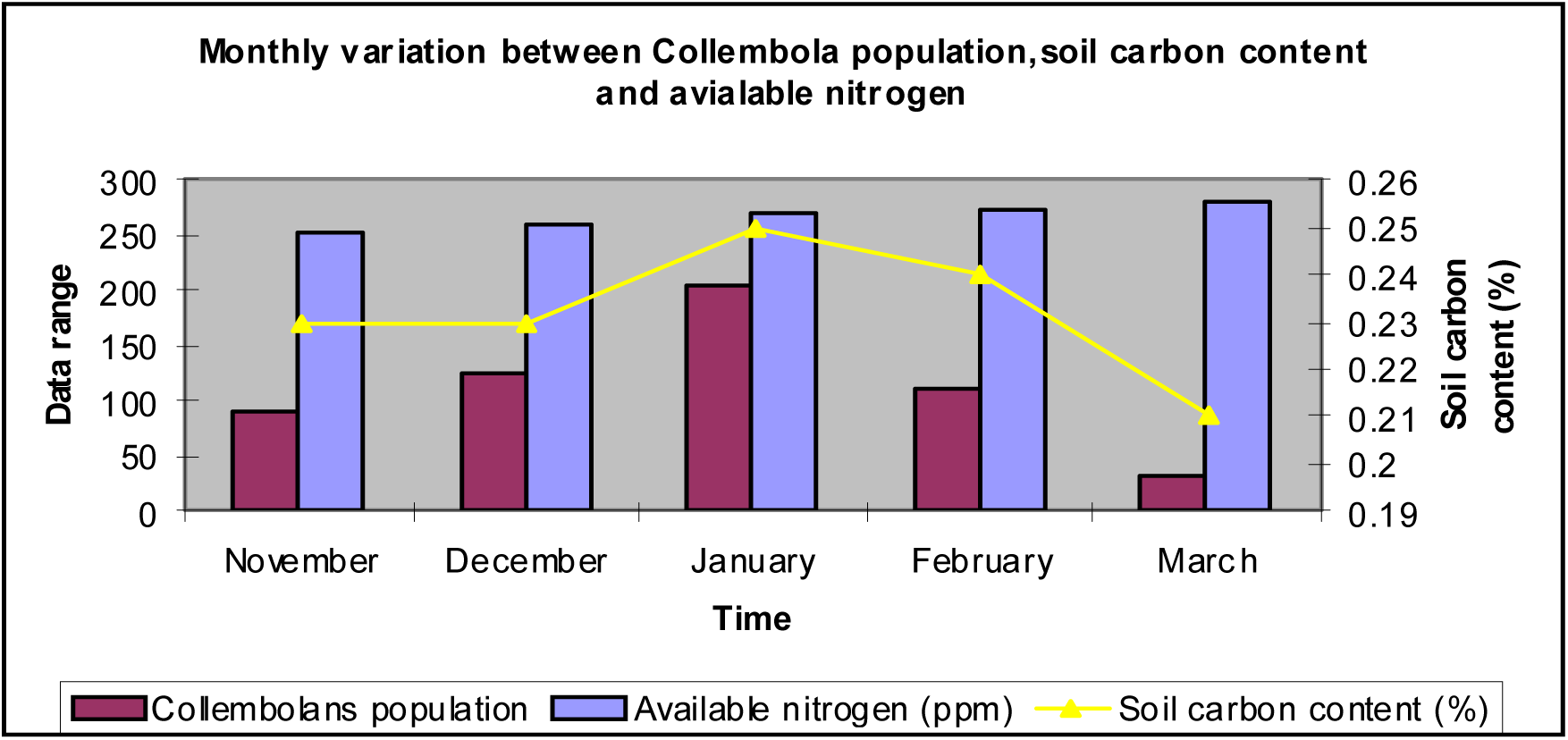
Monthly variation between Collembola population, soil carbon content and available nitrogen.

In this investigation, we observed that the less availability of carbon contents as well as higher concentration of nitrogen in soil, soil moisture regimes, temperature fluctuation in soil environment, absence or presence of food resources and intensive farming all have cumulative effect on collembolans diversity in agricultural soils that we observed in our study. The extra use of chemical fertilizers (fertilizer load) and mechanical tillage may also effect on the population of these natural currency. Thus, our results have similar trends that of Sonja et al. (1991), carried out a study and viewed that not a single factor but a cumulative action of a number of factors are responsible to control the seasonal periodicity of soil microarthropods population. It was reported in our study that, March was the least diversified month, may be due to the increased environmental temperature followed by relative humidity and decreased soil moisture contents in soil (table-2, figure-1).

In our study, the problem simply associated with taxonomy (identification) of soil microarthropods. Thus, numbers of soil microarthropod sampled do not indicate the actual numbers with in the entire community but simply represent an index of temporal distribution of soil microarthropod during investigation period. High population in winter months revealed that of cool temperature and medium humidity and approximately stable moisture content that was present in the soil (Table-2). In the present study, the real disturbance cannot be proved; however, density, abundance and survival of soil collembolans is still undergoes dynamic with reference to climate. Thus, the site specific productivity is a function of climate, soil properties and biotic potential of vegetation occupying the site (Knoepp et al. 2000). In the light of this sentence, we might expect that, wheat is a more diversified crop and occupying more floral diversity as well as root-root network. So, it can be stated that, high diversity of floras influence the soil microarthropods population. However, the clear cut evidence cannot be made on this aspect. Similarly, a huge degree of specific and temporal variation within a given land use/ecosystem was observed due to soil microbial properties and micronutrients by several other workers (Parkin 1993, Khan and Nortcliff 1982); However, our opinion is that, the contribution of a particular variable is an important task to a specific crop because, the species specific diversity within a particular crop may have profound affect on the nature and fertility of soil.

It is understood that the habitat management may also be feasible to maintain the key beneficial species in specific cropping system (Powell 1986) but this may not simply be applicable, because the interactions among the physical, chemical and biological components of agricultural system are inherently complex. Many workers have reported that the higher population buildup of soil microarthropods during rainy season and a sharp decline during summer months (Hazra and Sanyal 1996, Jam et al. 1986, Reddy et al. 1992); however peak population buildup of Collembola during winter months recorded in this study differs from majority of other reports. Only few studies have shown post monsoon increase or winter peak.

There has been considerable interest to arable the contribution of soil microarthropods specifically that of collembolans because they are sound functional attributes of soil quality as they are enormously diversified in native agricultural soils. However, the extreme climatic variations and their effect on soil microarthropods diversity are interesting in view of the great economic significance of Collembolans and its intimate relationship with climatic as well as seasonal variation. Thus, this investigation may be profitable in current knowledge of effect of climatic change and edaphic factors on the diversity of soil microarthropods specifically on collembolans in native agricultural soils. Under natural resource use and their conservation efforts, the findings of the study may also be profitable for researchers, farmers and policy makers.

This study has shown profound diversity in Collembolans population and highlights the significance of the variety of chemical and edaphic factors which regulate the fluctuation and diversity of microarthropods population in a varied manner. Thus, in a progressive approach, one would be expected to the significant and thoughtful experiment on seasonal variation of soil microarthropods in relation to weather patterns because, effect of climate change on the diversity and population dynamics of soil microarthropods specifically on collembolans can be a powerful tool to assess soil quality and soil health status.

## Conclusions

We investigate the diversity and relative abundance of Collembola in a wheat (*Triticum aestivum*) field at Aligarh. Following are the significant contributions of our study that-

- The species diversity of Collembola corresponds to Entomobryoidae, Isotomidae, Hypogastruridae and Sminthuridae in which, Hypogastrurides were dominant in entire community of Collembola.
- A close correlation of soil organic carbon contents with that of collembolans diversity was recorded in this study.
- The highest population of Collembolans recorded at neutral pH level.
- In terms of numbers of soil microarthropods, Collembola apparently constituted a better population than the other diverse group of soil microarthropods such as Acari(mites).
- The factors like soil moisture, temperature, relative humidity, soil pH, soil organic carbon contents as well as available nitrogen in the soil and food resources all have cumulative effect on the diversity of soil microarthropods.

